# Content bias in the cultural evolution of house finch song

**DOI:** 10.1101/2021.03.05.434109

**Authors:** Mason Youngblood, David Lahti

## Abstract

In this study, we used a longitudinal dataset of house finch (*Haemorhous mexicanus*) song recordings spanning four decades in the introduced eastern range to assess how individual-level cultural transmission mechanisms drive population-level changes in birdsong. First, we developed an agent-based model (available as a new R package called *TransmissionBias*) that simulates the cultural transmission of house finch song given different parameters related to transmission biases, or biases in social learning that modify the probability of adoption of particular cultural variants. Next, we used approximate Bayesian computation and machine learning to estimate what parameter values likely generated the temporal changes in diversity in our observed data. We found evidence that strong content bias, likely targeted towards syllable complexity, plays a central role in the cultural evolution of house finch song in western Long Island. Frequency and demonstrator biases appear to be neutral or absent. Additionally, we estimated that house finch song is transmitted with extremely high fidelity. Future studies should use our simulation framework to better understand how cultural transmission and population declines influence song diversity in wild populations.

## Introduction

Learned birdsong, which is socially transmitted and accumulates changes across generations, is a powerful model system for the study of cultural evolution [1–4]. Over the course of transmission, changes can accumulate through a variety of means, resulting in spatial and historical diversity in song structure [5, 6]. Transmission biases, or biases in social learning that predispose individuals to favor particular cultural variants, are important selective forces [7] that can result in significant changes at the population-level [8– 11]. Conformity bias (a bias for more common variants), for example, appears to play a key role in the cultural transmission of foraging techniques in great tits, and birdsong in swamp sparrows [12, 13].

In the last several decades, researchers have begun to explore how these transmission processes can be extracted from large-scale cultural datasets. This top-down approach, in which mechanisms of cultural transmission are inferred by looking at population-level changes in the variants themselves, was originally pioneered by archaeologists [14, 15] and has since been applied to various cultural domains including dog breeds [16], music [17], and birdsong [13, 18]. For example, chestnut-sided warblers have accented and unaccented songs that exhibit slightly different patterns of geographic variation [18]. By comparing their rates of turnover and mutation, researchers were able to conclude that only one of the two song types is subject to stabilizing selection, likely caused by some form of transmission bias [18]. In the current study we utilize a more recent version of this approach called generative inference, that allows researchers to test hypotheses related to specific forms of transmission bias [19].

Generative inference is a powerful simulation-based method that uses agent-based modeling and approximate Bayesian computation to infer underlying processes from observed data [19]. Agent-based modeling allows researchers to simulate generations of interacting individuals that culturally transmit information according to certain parameters, some of which are variable and some of which are constant, based on what is known about the system. By running many iterations of these simulations with different parameter values and comparing the output to the observed data, it is possible to infer the parameter values that likely generated the observed data [20–23]. Lachlan et al. [13] recently investigated the cultural transmission of swamp sparrow song using an agent-based model that incorporated three forms of transmission bias. They found that conformity bias best explained the cultural composition of their population [13]. For the current study, we modified this model to allow for dynamic population size. We used this model with a new version of approximate Bayesian computation that incorporates random forest machine learning [24] to infer whether transmission bias has played a role in the cultural evolution of house finch song.

The house finch is a socially monogamous, non-territorial passerine [25] with learned song [26] that culturally evolves [27] and is likely subject to sexual selection [28]. It is native to the western United States and Mexico, but was introduced from California to Long Island around 1940 [29, 30] and has since expanded throughout the eastern and central United States [31]. The house finch is an ideal model for such a study because we have a collection of songs from over a thousand individuals from an extensive geographical area in the introduced eastern range recorded between 1971–1975 [32], 2010–2016 [33], and 2018–2019. Our recent comparative study of house finch songs recorded in 1975 and 2012 in Long Island showed that local dialects have become less distinct and both song and syllable diversity have increased at the population-level [33]. Additionally, although none of the songs from 1975 have persisted, around half of the syllable types recur in 2012 [33]. We proposed three possible drivers for the increase in diversity: (1) higher innovation rates, (2) more demonstrators (as a result of increased density), and (3) more dispersal. Although the average dispersal distance and number of demonstrators probably increased during the population expansion of the 1980s, these variables would also have been affected by the significant bottleneck that occurred in the 1990s as a result of mycoplasmal conjunctivitis [34]. In addition, it is unclear, with the methods used, whether or not the increase in diversity is the result of neutral or biased cultural transmission.

The aim of the this study was to infer whether transmission bias played a role in the cultural evolution of house finch song during a rapid demographic expansion and crash in the introduced eastern range. We investigated the three forms of cultural transmission bias, first described by Boyd and Richerson [35], that likely play a role in birdsong [13]: content bias, frequency bias, and demonstrator bias. Content bias occurs when some cultural variants are more likely to be learned because of the content of those variants (e.g. frequency bandwidth and complexity) [36]. The most widespread form of content bias in birds is a preference for species-typical song [4, 37]. In white-crowned sparrows, for example, birds are more likely to learn phrases from songs that contain a species-specific introductory whistle [38]. Content biases can be adaptive, given that females choose mates based on particular acoustic features [39, 40]. Since female house finches prefer elaborate song [28, 39], and complex syllables are overrepresented in our more recent recordings [33], content bias, specifically for syllable complexity, may be present in male house finches.

Frequency biases (e.g. conformity bias and novelty bias) occur when the commonness or rarity of cultural variants affects their adoption [36]. In birds, conformity is the most well-documented form of frequency bias [4, 13, 41, 42]. White-crowned sparrows, for example, selectively retain the songs of their neighbors after establishing territories, resulting in stable local dialects [43]. Under certain conditions, conformity bias is thought to be broadly adaptive because it allows individuals to quickly adopt common variants [35, 44], and is particularly important in birds because females of some species prefer males who sing local songs [45, 46]. That being said, the adaptive nature of conformity bias depends on several assumptions that may not hold for all systems [47], and weak conformity is likely to be more adaptive than strong conformity in many contexts because it allows potentially advantageous innovations to spread [48]. To our knowledge, the only clear evidence of novelty bias in birds is in medium ground finches. In this species, males appear to preferentially learn rarer songs, and those that sing rarer songs survive longer and produce more offspring [49]. Gibbs attributes this novelty bias to the high population density of territorial males, a situation in which birds with more distinct songs may spend less time displaying and more time foraging and avoiding predation [49]. Other possible examples of novelty bias, such as song “revolutions” in corn buntings and continent-level shifts in white-throated sparrow songs, [50, 51], could also result from content bias and need to be investigated further. In house finches, a non-territorial species in which females prefer males who sing local songs [52], conformity bias is more likely to be adaptive than novelty bias.

Demonstrator bias occurs when cultural variants produced by particular demonstrators are preferred [36]. In indigo buntings, for example, young birds preferentially learn songs produced by males that they socially interact with [53]. In other species, birds are more likely to learn from males that are older [41] or more aggressive [54]. In the house finch, a species in which both song and redness are used in mate choice [39], it is possible that young birds pay more attention to redder demonstrators. Given that social context influences song production [55], demonstrator bias in house finches could also be the result of observational learning [56], in which young birds pay attention to which males spend the most time with females and choose demonstrators accordingly.

One of the biggest challenges in extracting transmission biases from population-level data is dealing with equifinality, or the fact that different transmission mechanisms can yield similar results [57]. This is particularly relevant for this study, as content, frequency, and demonstrator biases can yield similar population-level patterns [58]. For example, a content bias for songs containing particular syllables, such as the introductory whistle in white-crowned sparrows [38], could generate stable dialects that are consistent with conformity bias. Alternatively, a content bias for complex syllables that only the highest-quality males can produce could manifest as novelty bias. However, despite potential similarities in emergent pattern, simulations indicate that content, frequency, and demonstrator biases have discriminable effects at the population-level [13]. Content bias, for example, tends to reduce the number of rare variants in the population by increasing the turnover of unattractive new variants. Demonstrator bias, on the other hand, tends to cause a decrease in cultural diversity by reducing the number of potential demonstrators [13]. By simulating all three processes simultaneously, we aim to infer which is dominant in the cultural evolution of house finch song.

## Methods

### 0.1 Recording

We used recordings from 1975 [32], 2012 [33], and 2019 that were collected in western Long Island (Brooklyn, Queens, and Nassau County) (Figure 1). The 1975 and 2012 recordings were identical to those recently analyzed by Ju et al. [33], whereas the 2019 recordings were specifically collected for this study. Recordings from 1975 were collected with a Nagra III reel-to-reel tape recorder and a Sennheiser 804 shotgun microphone, and converted to digital files (32-bit, 96 kHz) by the Cornell Lab of Ornithology in 2013 [33]. Recordings from 2012 (16-bit, 44.1 kHz) and 2019 (32-bit, 48 kHz) were collected with a Marantz PD661 solid state recorder and a Sennheiser ME66 shotgun microphone. The recordings from 1975 were downsampled to 48 kHz prior to analysis (Luscinia processes both 44.1 kHz and 48 kHz files). In all three years special precautions were taken to avoid recording the same bird twice [32, 33]. Each site was visited only once. Within a site, only one individual was recorded within a 160 m radius until they stopped singing or flew away.

**Figure 1.**
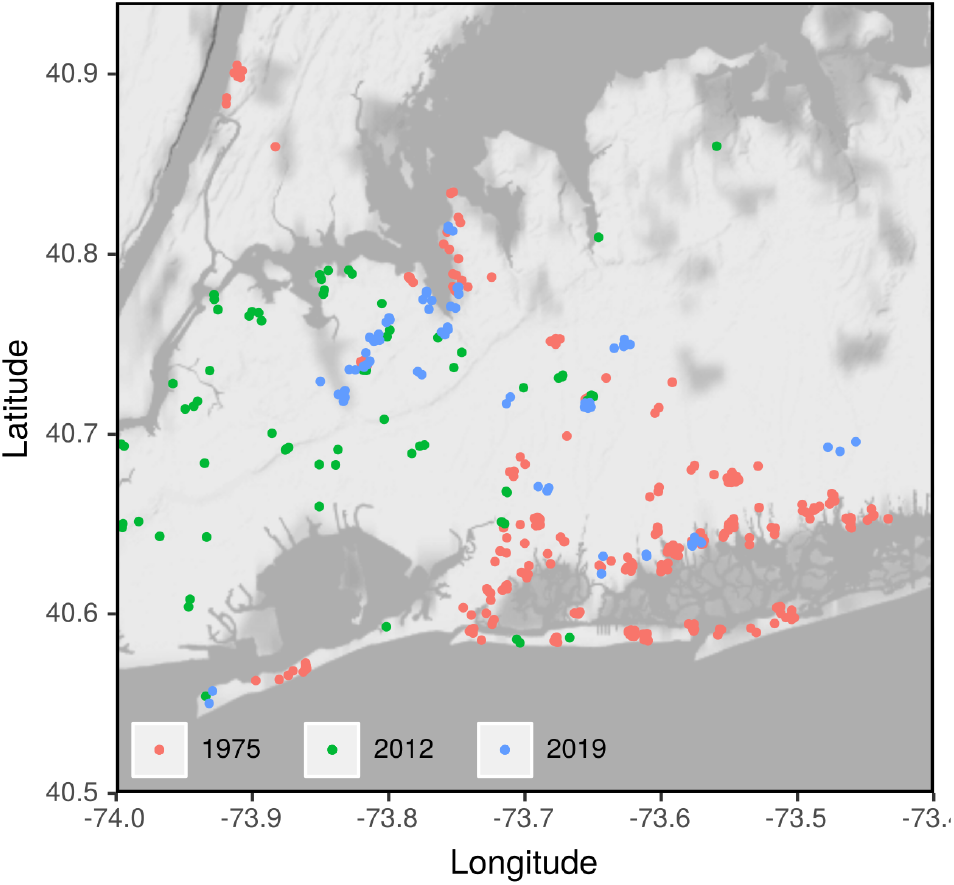
Locations of recordings from 1975, 2012, and 2019 in western Long Island (Brooklyn, Queens, and Nassau County).

### 0.2 Song Analysis & Clustering

Song analysis was conducted in Luscinia (http://rflachlan.github.io/Luscinia/), a database and analysis program developed specifically for birdsong. Songs were analyzed with a high pass threshold of 2,000 Hz, a maximum frequency of 9,000 Hz, and 5 dB of noise removal. We excluded 965 songs (26.2%) from the analysis due to high levels of noise or overlap with other bird species. Continuous traces with more than 20 ms between them were classified as syllables [32]. Some syllables were further separable into elements, or traces with less than 20 ms between them. See Figure 2 for an example of an analyzed song. Cluster analysis could not be done in Luscinia due to computational limitations, so we exported the raw mean frequency measurements and analyzed them in R. Mean frequency here refers to the mean frequency in each 1 ms bin, so it is a continuous trace across the length of syllables. Mean frequency was log-transformed prior to analysis [13]. First, the normalized distances between all of the syllables were calculated via dynamic time warping (DTW) with a window size of 10 (10% of the average signal length) using the *dtwclust* package [59]. A window size of 10% of the signal length is commonly used in speech processing research and seems to be a practical upper limit for many applications [60]. Infinite distances (0.19%) caused by comparisons of syllables with extreme signal length differences were assigned the maximum observed distance value. Next, hierarchical clustering and dynamic tree cut were used to cluster the syllables into types [33, 61]. Hierarchical clustering was conducted with the UPGMA method implemented in the *fastcluster* package [62], and dynamic tree cut was conducted with the *dynamicTreeCut* package [63]. For dynamic tree cut, we ran the hybrid algorithm with a minimum cluster size of 1 to maximize the representation of rare syllable types. We chose a deep split value of 3 because it resulted in relatively good cluster validity indices (see Table S1) and a similar number of syllable types compared with previous studies of house finch song [33, 61].

**Figure 2.**
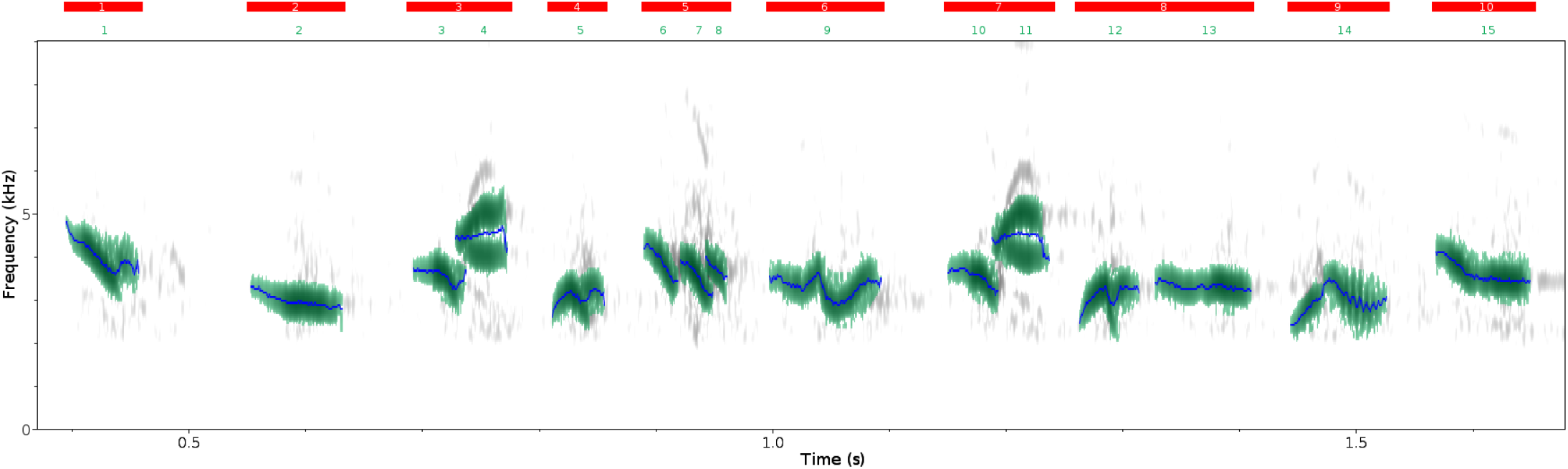
The first 10 syllables from a song recorded in 1975 and analyzed in Luscinia. Each red bar corresponds to a syllable, and each green number corresponds to an element within that syllable. The blue traces represent mean frequency. In this song, syllable three and syllable eight were classified as the same syllable type during dynamic time warping, and all other syllables are unique types.

In order to control for variation in repertoire coverage, we only included birds from whom we recorded enough songs to cover at least 75% of their syllable repertoire [33]. To do this, we plotted the cumulative proportion of syllables types captured with each additional song for the 106 individuals with at least 10 recorded songs as per Ju et al. [33] (see Figure S3). On average, 78.0% of an individual’s repertoire was captured in the first eight songs, so we only calculated repertoire sizes and observed summary statistics from birds with at least eight recorded songs.

### 0.3 Simulation & Generative Inference

The agent-based model was adapted from Lachlan et al. [13] and written in R and C++. For computational efficiency, we only included males in our model. Female house finches rarely sing in the wild [64], and thus are unlikely to act as demonstrators to juvenile birds. The model of cultural transmission is initialized with *N*_*B*_ birds and *N*_*S*_ possible syllables. Repertoire sizes are drawn randomly from a normal distribution based on the observed data. Each bird is assigned a demonstrator attractiveness index (*t*_*m*_), calculated by raising Euler’s number to a random number drawn from a normal distribution with a mean of 0 and a standard deviation of *v* (so that the indices are non-negative and centered around 1). Each bird is also assigned a random geographic index (*g*) between 1 and 100, which is used to simulate geographic location. Each syllable is assigned an attractiveness index (*M*) of 1 (attractive) or 0.05 (unattractive), where *p*_*att*_ is the proportion of all syllables that are attractive. At the start of each year a new generation learns syllables from a pseudorandom set of *D* demonstrators, where the probability of a bird learning from a demonstrator is the inverse of the absolute difference between their geographic indices (i.e. more geographically distant adults are less likely to act as demonstrators). Dispersal rates in western Long Island can be as high as 91.3% [65] and males appear to learn most of their repertoire after dispersal [64], indicating that choosing demonstrators pseudorandomly by geographic location after dispersal is appropriate.

The probability of a bird learning a syllable (*P*_*x*_) depends upon the frequency of that syllable among the demonstrators (*F*_*x*_), the attractiveness of the syllable (*M*_*x*_), and the mean attractiveness of the demonstrators singing the syllable (*T*_*x*_). To simulate frequency-based bias, *F*_*x*_ is raised to the exponent *α*, where *α >* 1 corresponds to conformity bias and *α <* 1 corresponds to novelty bias. As such, 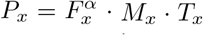. During the learning process, new syllable types are innovated with probability *µ*. We use the term innovation to describe the generation of new variants, as is common in both animal and human cultural evolution research [4, 66], but our model is agnostic about the exact processes involved (copy error, guided variation, etc.). Innovated syllables are drawn from the initial pool of possible syllables (*N*_*S*_), to reflect the production limitations imposed by species-specific physiological and cognitive constraints [26, 67]. Mortality is simulated at the end of each year by randomly selecting 50% of the agents to remove (overwinter survival varies from 39-63% in different populations [68–70]), and at the beginning of each year new individuals are added to bring the population up to size.

In summary, *p*_*att*_, *α*, and *v* correspond to content, frequency, and demonstrator bias, respectively. Smaller values of *p*_*att*_ are indicative of stronger content bias, as fewer syllables are attractive to learners. Values of *α >* 1 are indicative of conformity bias, while values of *α <* 1 are indicative of novelty bias. And lastly, higher values of *v* are indicative of of stronger demonstrator bias, as there is more variation in demonstrator attractiveness.

To model changes in population size, we extrapolated from *N*_*B*_ using data from the Christmas Bird Count (CBC) (https://www.audubon.org/conservation/science/christmas-bird-count). Annual trends were estimated by calculating the percent change in the average number of house finches encountered per party hour in Brooklyn, Queens, and Nassau County for each year. For example, if the simulated population size in year *x* was 100 and the CBC indicated a 25% increase between *x* and *x* + 1, then the population size in year *x* + 1 would be 125. These annual trends from the CBC can be seen in Figure S1. The resulting population estimates were halved in the model, since only males were included.

Following an initial burn-in phase, each iteration of the model was run from 1970 to 2019. A 100 year burn-in phase appears to be sufficient to reach an equilibrium-level of diversity (Figure S4). In 1975, 2012, and 2019 the following summary statistics were calculated from a random sample of the same size as the corresponding observed sample: (1) the proportion of syllables that only appear once, (2) the proportion of the most common syllable type, (3) the number of syllable types, (4) Simpson’s diversity index, (5) Shannon’s diversity index, (6) Pielou’s evenness index, and (7) the exponent of the fitted power-law function to the progeny distribution [13, 21]. We chose these summary statistics because they capture diversity with an emphasis on both common and rare variants, as well as the general shape of the frequency distribution. The three sets of seven summary statistics from each simulation were compared with the same summary statistics from the same years in the observed dataset during parameter estimation.

Parameter estimation was conducted with the random forest algorithm of approximate Bayesian computation, using the *abcrf* package in R [24]. Random forest is a form of machine learning in which a set of decision trees are trained on bootstrap samples of variables, and used to predict an outcome given certain predictors [71]. Traditional approximate Bayesian computation methods function optimally with fewer summary statistics, requiring researchers to reduce the dimensionality of their data [72]. We chose to use random forest for parameter estimation because it appears to be robust to the number of summary statistics, and does not require the exclusion of potentially informative variables [73]. In addition, it requires fewer iterations of the model compared to other methods, making it much less computationally intensive [73]. Random forest approximate Bayesian computation was conducted with the following steps:

1. 200,000 iterations of the model were run to generate simulated summary statistics for different values of the parameters: *N*_*B*_, *N*_*S*_, *µ, D, p*_*att*_, *α*, and *v*.
2. The output of these simulations was combined into a reference table with the simulated summary statistics as predictor variables, and the parameter values as outcome variables.
3. A random forest of 1,000 regression trees was constructed for each of the seven parameters using bootstrap samples from the reference table. A maximum tree depth of five was used to avoid overfitting.
4. Each trained forest was provided with the observed summary statistics, and each regression tree was used to predict the parameter values that likely generated the data.

The following parameters were assigned log-uniform prior distributions, so that more sampling occurred near the lower bounds: *N*_*S*_ = {596–800}, *µ* = {0.001–0.3 }, *v* = {0.01–6 },and *D* = 2–10 (modified from Lachlan et al. [13]). Increased sampling near the lower bounds improves the resolution of posterior distributions for parameters that are more likely to have low or moderate values [13]. The lower bound of *N*_*S*_ is set to 596 which is the number of syllables identified in our dataset. The following parameters were assigned uniform prior distributions, so that sampling was unbiased across the entire range: *N*_*B*_ = 2,000–10,000 *p*_*att*_ = {0.01–1} and *α* = {0.25–3 }(modified from Lachlan et al. [13]). Assuming an average population density of 3.8 birds/km^2^ [74, 75], our estimate for the average population size of western Long Island (1,203.4 km^2^) is 4,573. We chose The posterior distributions for *p att, α*, and *v* can be seen a wide uniform prior distribution for initial population size that includes this estimate but allows for significant variation over time.

### 0.4 Logistic Regression

To assess whether syllables with particular characteristics had higher persistence, we conducted Bayesian logistic regression with all of the syllable types detected in 1975 using the *rstanarm* package [76]. We started out with the following predictor variables used by Ju et al. [33]: average frequency (Hz), minimum frequency (Hz), maximum frequency (Hz), bandwidth (Hz), duration (ms), concavity (changes in the sign of the slope of the mean frequency trace per ms), and excursion (cumulative absolute change in Hz per ms). Concavity and excursion are both indicators of syllable complexity [33, 77]. Concavity was calculated after smoothing the mean frequency trace using a polynomial spline with a smoothing parameter of 5 (chosen by visual inspection). The predictor variables were averaged across all of the observations of each syllable type. Maximum frequency, average frequency, and bandwidth were left out of the modeling due to multicollinearity issues (*V IF >* 4). Whether or not the syllable type persisted to 2019 (logical: T/F) was used as the outcome variable. We used Student’s *t*-distributions with a scale of 2.5 as priors to allow for a relatively wide range of parameter estimates.

## Results

The dynamic tree cut identified 596 syllables types across the 2,724 songs from 331 individuals that we analyzed. After applying the eight song threshold to ensure that we had at least 75% of individuals’ syllable repertoires, we calculated the observed summary statistics from 36, 75, and 29 individuals from 1975, 2012, and 2019, respectively (Table S2). The observed mean repertoire size of 61.16 (*SD* = 17.55)was used for all of our simulations. in Figure 3 and Table 2. The median estimate for *p*_*att*_ was 0.21, with a 95% CI that falls far short of 1. This indicates that the cultural changes in our dataset are consistent with strong content bias. The median estimate for *α* was 0.80 which is consistent with moderate novelty bias, but as the posterior distribution peaks at 1 and the 95% CI overlaps 1 we cannot reject the null hypothesis. The 95% CI for *v* encompasses almost all of the prior distribution, and the out-of-bag error for the random forest is orders of magnitude higher than the other parameters. This indicates that demonstrator bias is either not present or does not influence cultural changes at the population-level. The 10 most important summary statistics used by the random forests to estimate each parameter, identified using the Gini impurity method, can be seen in Table S3.

**Figure 3.**
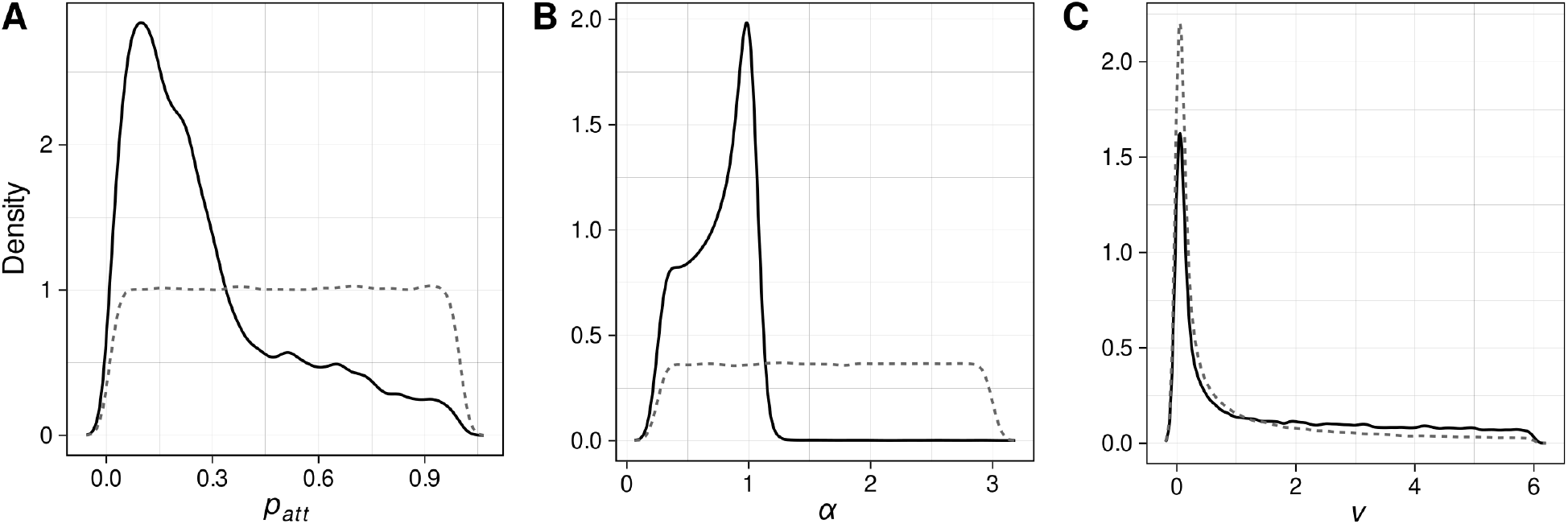
The prior (dotted lines) and posterior (solid lines) distributions for the three dynamic parameters in the agent-based model related to transmission biases. *p*_*att*_, *α*, and *v* correspond to content, frequency, and demonstrator bias, respectively.

**Table 1:** The description and prior distribution for each dynamic parameter in the agent-based model.

|  | *Description* | *Prior* | *Type* |
| --- | --- | --- | --- |
| $N_B$ | Initial population size | {2,000–10,000} | Uniform |
| $N_S$ | Total number of syllables | {596–800} | Log-uniform |
| $\mu$ | Innovation rate | {0.001–0.3} | Log-uniform |
| $D$ | Number of demonstrators | {2–10} | Log-uniform |
| $p_{att}$ | Proportion of attractive syllables | {0.01–1} | Uniform |
| $\alpha$ | Level of conformity bias | {0.25–3} | Uniform |
| $v$ | Variation in demonstrator attractiveness | {0.01–6} | Log-uniform |

**Table 2:** The median, 95% credible interval, and out-of-bag normalized mean absolute error of the posterior distribution for each dynamic parameter in the agent-based model.

|  | *M* | *95% CI* | *NMAE* |
| --- | --- | --- | --- |
| $N_B$ | 5830 | [2174, 9789] | 0.46 |
| $N_S$ | 635 | [598, 777] | 0.045 |
| $\mu$ | 0.0067 | [0.0011, 0.17] | 1.55 |
| $D$ | 4 | [2, 10] | 0.51 |
| $p_{att}$ | 0.21 | [0.023, 0.89] | 0.64 |
| $\alpha$ | 0.80 | [0.28, 1.06] | 0.35 |
| $v$ | 0.73 | [0.012, 5.64] | 16.55 |

The posterior distributions for the other parameters can be seen in Figure S5 and Table 2. The 95% CIs for *N*_*B*_ and *D* are nearly equivalent to the prior distributions, indicating that neither has a significant effect on the cultural changes in our dataset. *N*_*S*_ had a median estimate of 635 and a broad 95% CI that excludes values above 777. This means that, at the current resolution of the hierarchical clustering, we captured at least 76.71% of the possible syllable types in the population. Lastly, the median estimate for *µ* is 0.0067 with a 95% CI that excludes values above 0.17, suggesting that house finches culturally transmit song with high fidelity like swamp sparrows [13]. This result should be interpreted with caution, as *µ* may be underestimated by the log-uniform prior distribution, and the out-of-bag error is relatively high compared to other parameters.

When both *α* and *v* are near neutrality (0.99 *< α <* 1.01, *v <* 0.02), there are significant positive correlations between *p*_*att*_ and the proportion of syllables that only appear once (*R* = 0.17, *p <* 0.05), the number of syllable types (*R* = 0.84, *p <* 0.0001), Simpson’s diversity index (*R* = 0.69, *p <* 0.0001), Shannon’s diversity index (*R* = 0.86, *p <* 0.0001), Pielou’s evenness index (*R* = 0.52, *p <* 0.0001), and the exponent of the fitted power-law function (*R* = 0.22, *p <* 0.05), and there is a significant negative correlation between *p*_*att*_ and the proportion of the most common syllable type (*R* = *-*0.69, *p <* 0.001). This indicates that stronger content bias (lower value of *p*_*att*_) reduces cultural diversity in favor of attractive syllable types independently of other factors, which may explain why we were unable to get strong estimates for parameters like population size and the number of demonstrators.

The results of the Bayesian logistic regression are shown in Table 3. The persistence of syllable types from 1975 to 2019 is positively predicted by both minimum frequency and concavity, and is negatively predicted by duration. Excursion does not significantly predict syllable persistence.

**Table 3:** The median, 95% credible interval, and standard deviation for each parameter included in the Bayesian logistic regression. Concavity is the number of changes in the sign of the slope of the mean frequency trace per ms, and excursion is the cumulative absolute change in Hz per ms. 95% credible intervals that do not overlap with 0 indicate a statistically significant effect.

|  | *M* | *95% CI* | *SD* |
| --- | --- | --- | --- |
| Min frequency (Hz) | 0.54 | [0.20, 0.93] | 0.18 |
| Duration (ms) | -1.34 | [-1.75, -0.97] | 0.21 |
| Concavity | 0.56 | [0.27, 0.89] | 0.16 |
| Excursion | -0.26 | [-0.53, 0.02] | 0.14 |

## Discussion

By applying simulations and machine learning to three years of birdsongs spanning four decades, we have provided evidence that content bias plays a central role in the cultural evolution of house finch song in western Long Island. Syllables with higher concavity, or syllables with more transitions in slope, were more likely to persist between 1975 and 2019. As such, we hypothesize that the strong content bias observed in this study is targeted towards syllable complexity. This is roughly consistent with previous research that found that syllables with higher excursion (another indicator of complexity) were more prevalent in individuals’ repertoires in 2012 [33], although this variable was not significant in our model. In house finches, courtship songs are longer and contain longer syllables than solo songs [55], and males that sing longer songs at a faster rate are more attractive [28] and have higher reproductive performance [39]. If female preferences for elaborate and energetically-costly songs extend to the syllable level, then males with a content bias for syllable complexity will have higher reproductive success.

Additionally, we estimated that house finch song is transmitted with a median fidelity of 99.99% and a lower bound (5% quantile) of 83%. This is consistent with the high fidelity of swamp sparrow song learning estimated by Lachlan et al. [13]. Future studies should investigate whether departures from perfect fidelity in house finches are directional and maximize vocal performance (e.g. increased complexity). For example, swamp sparrows accurately copy attractive tutors and speed up the trill rates of slower tutors [78]. House finches shorten and increase the frequency of canary trills learned in captivity [26] and may also modify conspecific syllables in non-random ways.

Interestingly, we did not find evidence for conformity bias, which appears to be relatively common in birds [4, 41, 42] and was recently identified in swamp sparrows using similar methods [13]. A previous study that found that female house finches prefer local over foreign songs, which is often associated with selection for conformity bias in males [45], tested birds from the introduced range (Ontario) with songs from the native range (Arizona) [52]. Preference differences between such distant and ecologically distinctive locations may be more reflective of morphological divergence (e.g. traits like bill depth [70, 79]) than regional song variation relevant for mating. Additionally, female preferences for local song in the same study appeared to be genetically inherited rather than learned [52]. If song preferences in both females and males are independent of early experience then frequency and demonstrator biases are less plausible in house finches.

We also found that syllables with a higher minimum frequency and a lower duration were more likely to persist between 1975 and 2019. A similar effect of minimum frequency on syllable persistence was observed by Ju et al. [33], who attributed it to the effect that low-frequency urban noise has on birdsong. If syllables with lower minimum frequencies are more likely to be masked by urban noise or less likely to be sung by males in noisy conditions, then they will be less likely to persist into the next generation [80]. Urban noise has been associated with increased minimum frequency in other house finch populations [81], and appears to be a plastic behavioral change on the part of males [82, 83]. The negative effect of duration on syllable persistence is more puzzling, as male house finches in noisy environments shorten some song types and extend some syllable types [83]. It is also unlikely to be due to content bias, as males appear to sing longer syllables in courtship songs [55]. One possibility is that short syllables resulting from homoplasy are less distinctive and more likely to be lumped together during hierarchical clustering, causing their persistence to be overestimated by the logistic regression.

There are several limitations to this study that should be highlighted. Firstly, our model is largely agnostic about guided variation and other internal transformative processes. Setting a maximum number of possible syllables to reflect physiological constraints may have indirectly captured some of these processes, but researchers applying these methods in more complex and domain-general cultural systems should carefully consider this issue. Secondly, our clustering process had a subjective component, in that we chose parameter values that yielded results consistent with previous studies. Since hierarchical clustering leads to a nested structure different parameter values for the tree cut just change the scale of the data, and the transmission biases we modeled should lead to similar population-level patterns regardless of scale. Finally, physiological and cognitive constraints likely limit the phonological space in which house finches can innovate, leading to widespread homoplasy over long periods of time [61] that makes it difficult to reconstruct the evolutionary history of syllable types [33]. While we explicitly accounted for homoplasy in our simulations by pulling innovated syllables from a limited number of possible types, we cannot rule out the fact that it may have influenced the results of our logistic regression.

The importance of cultural traits in conservation is increasingly recognized among scientists [84–88] and policymakers [89]. For songbirds that culturally transmit mating signals, researchers are concerned about the negative effects of population decline and habitat fragmentation on song diversity [90–99]. Reduced song diversity has been linked to decreased population growth [100], leading some researchers to suggest that song diversity may be a useful assessment tool for threatened populations of songbirds and cetaceans [101–104]. In order to use song diversity as a metric for population viability, researchers need to better understand how population diversity results from individual-level processes such as innovation and biased transmission [105]. An over-reliance on conformity bias, for example, could slow recovery by reducing effective variation, whereas novelty bias could quickly spread innovated syllables through the population [106]. If researchers using song diversity for population assessment assume that cultural transmission is neutral, which is unlikely to be realistic in most cases [106], they may underestimate long-term population viability. Future studies should use our agent-based modeling framework to better understand how cultural transmission and population declines influence song diversity in wild populations.

## Acknowledgments

We would like to thank Paul Mundinger and Frances Geller for recording in 1975 and 2012 (respectively), Sheila Gogineni and Christina Takos for recording in 2019, and Shari Zimmerman, Andrea Lopez, Ratna Kanhai, and Joderick Castillo for analyzing the songs in Luscinia. This research was supported, in part, under National Science Foundation Grants CNS-0958379, CNS-0855217, ACI-1126113 and the City University of New York High Performance Computing Center at the College of Staten Island.

## Data & Code Availability

The agent-based model used in this study is available as a new R package called *TransmissionBias* on Github (https://github.com/masonyoungblood/TransmissionBias). The “example” subfolder of the R package includes our processed syllable data and analysis scripts. The unprocessed song recordings are available upon request.

## Supporting information

**Figure S1:**
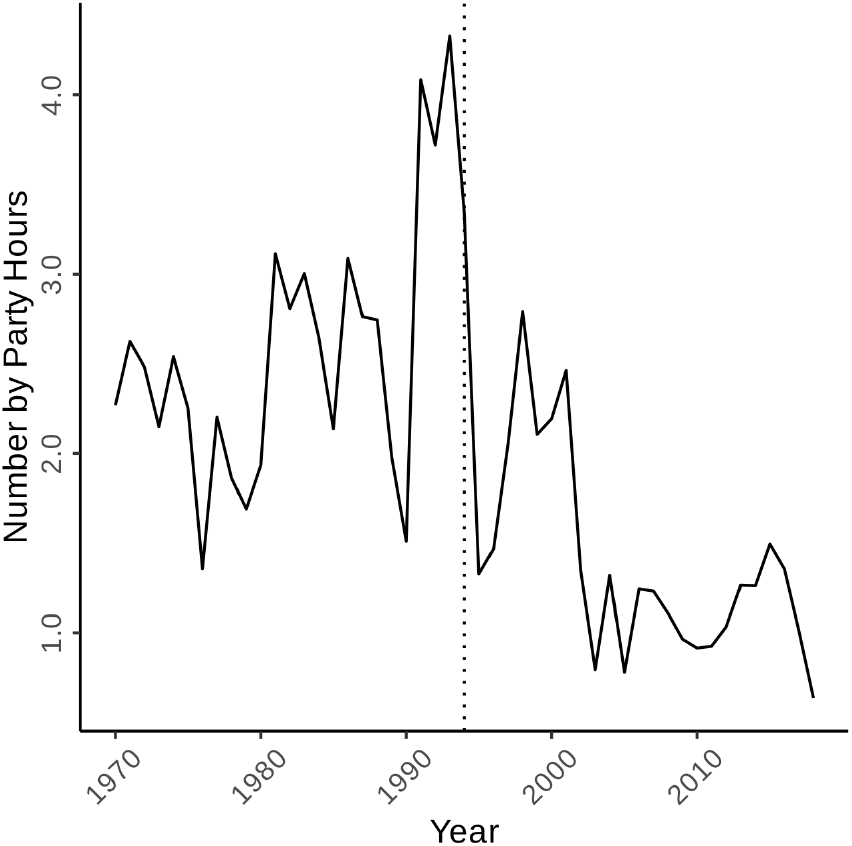
Average number of house finches encountered per party hour in western Long Island (Brooklyn, Queens, and Nassau County). The dotted line indicates 1994, the year in which conjunctivitis was first detected in the eastern range (Dhondt et al., 1998).

**Figure S2:**
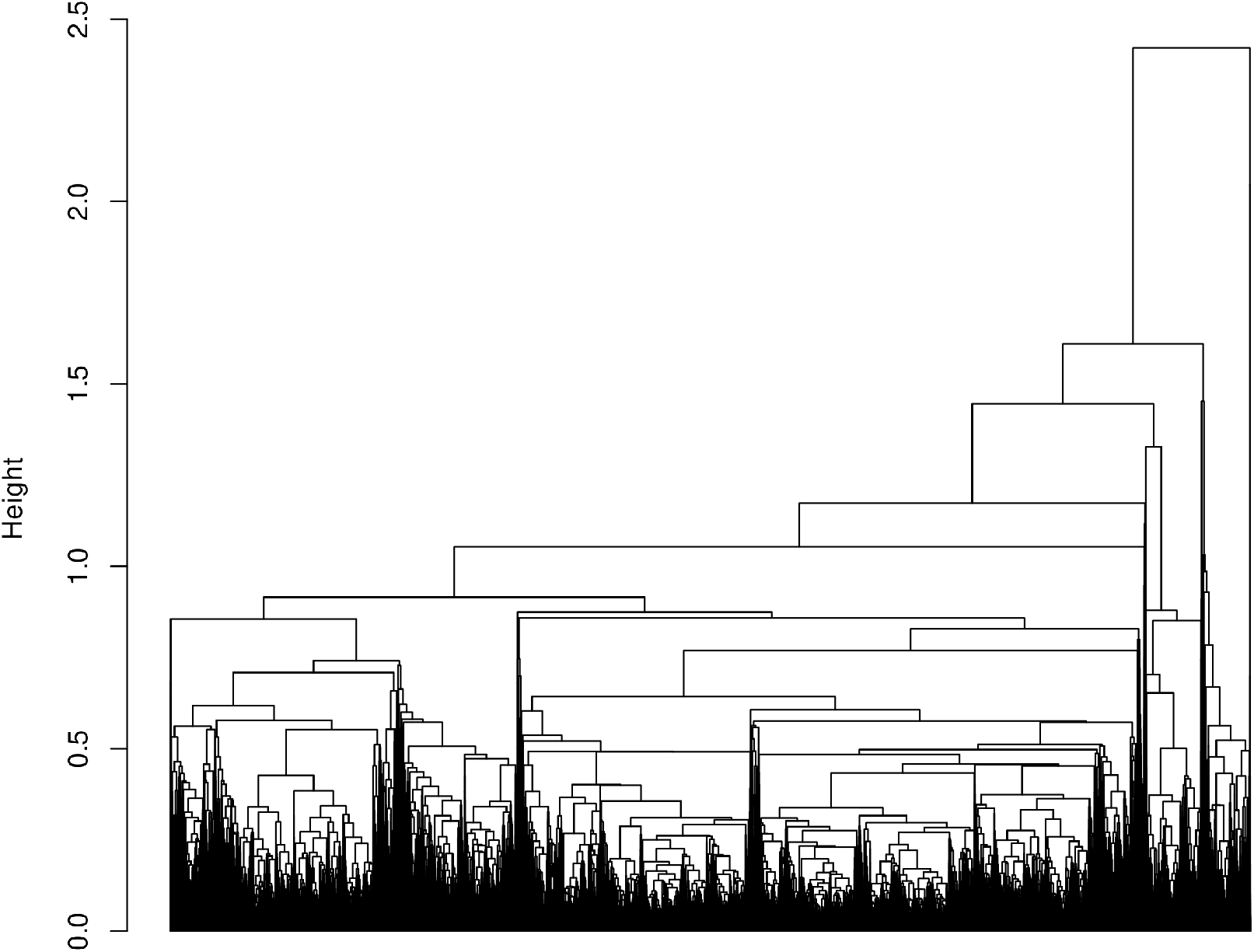
A dendrogram showing the results of the hierarchical clustering. The clusters could not be labelled because the number of syllable types exceeds the maximum color limit of R.

**Table S1:** The number of syllable types and cluster validity indices for each value of deep split, calculated using the *clValid* package [1]. For all three indices higher values correspond to higher quality clustering. We used a deep split value of 3 because it yielded results more consistent with previous research and had only slightly lower values of the silhouette and connectivity indices and an identical Dunn index, which appears to outperform the silhouette index when hierarchical clustering is used with real datasets [2].

| *Deep split* | *N* | *Silhouette* | *Dunn* | *Connectivity* |
| --- | --- | --- | --- | --- |
| 0 | 13 | 0.093 | 0.0027 | 7,168 |
| 1 | 36 | -0.030 | 0.0023 | 12,380 |
| 2 | 114 | 0.018 | 0.0020 | 20,797 |
| 3 | 596 | 0.071 | 0.0020 | 34,602 |
| 4 | 1,646 | 0.084 | 0.0020 | 44,967 |

**Figure S3:**
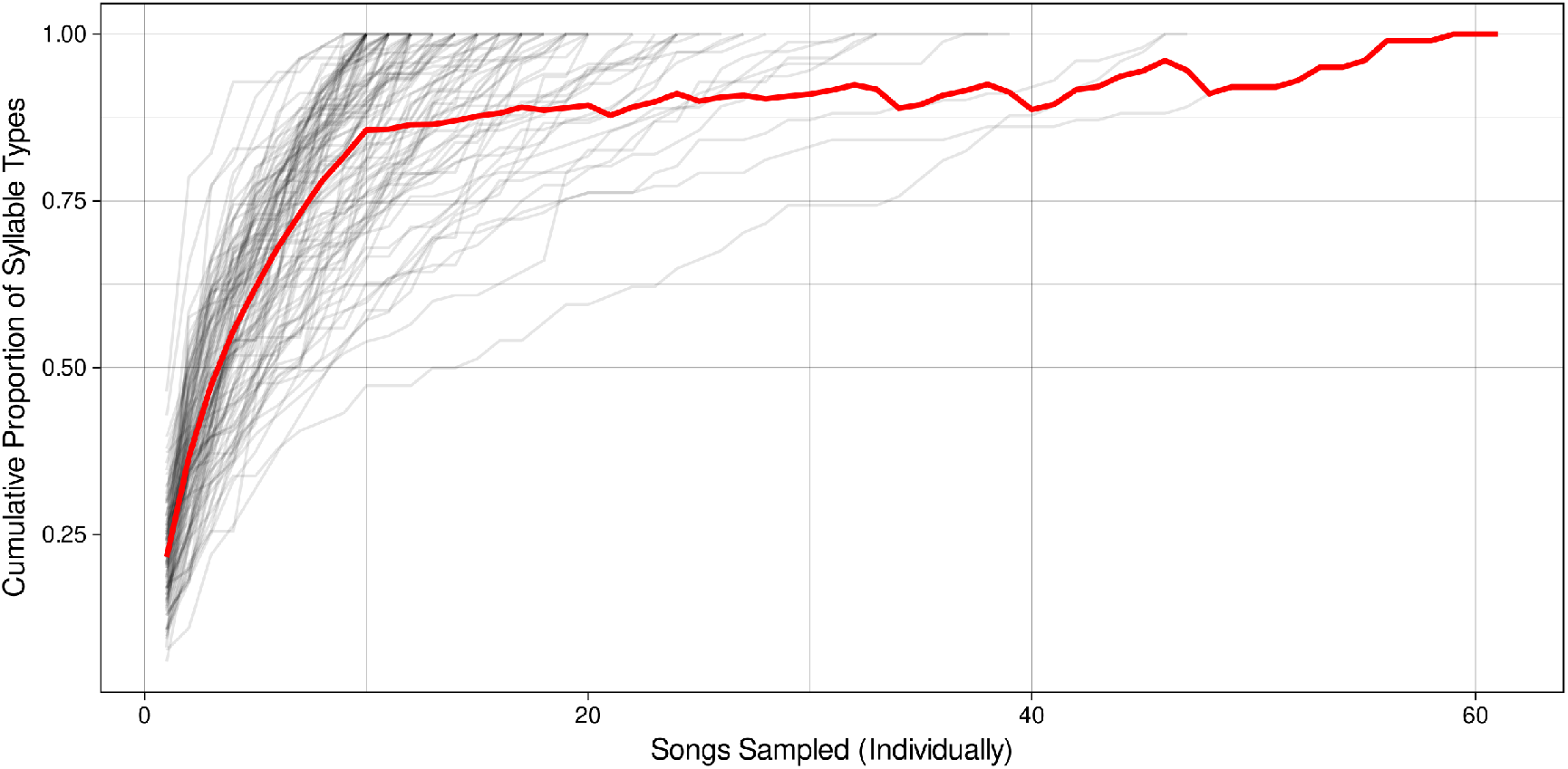
The cumulative proportion of syllable types (*y*-axis) detected with each additional song sampled (*x* -axis) for the 106 birds from whom we have at least 10 recorded songs. The red line is the average value, which cross 75% around eight songs.

**Figure S4:**
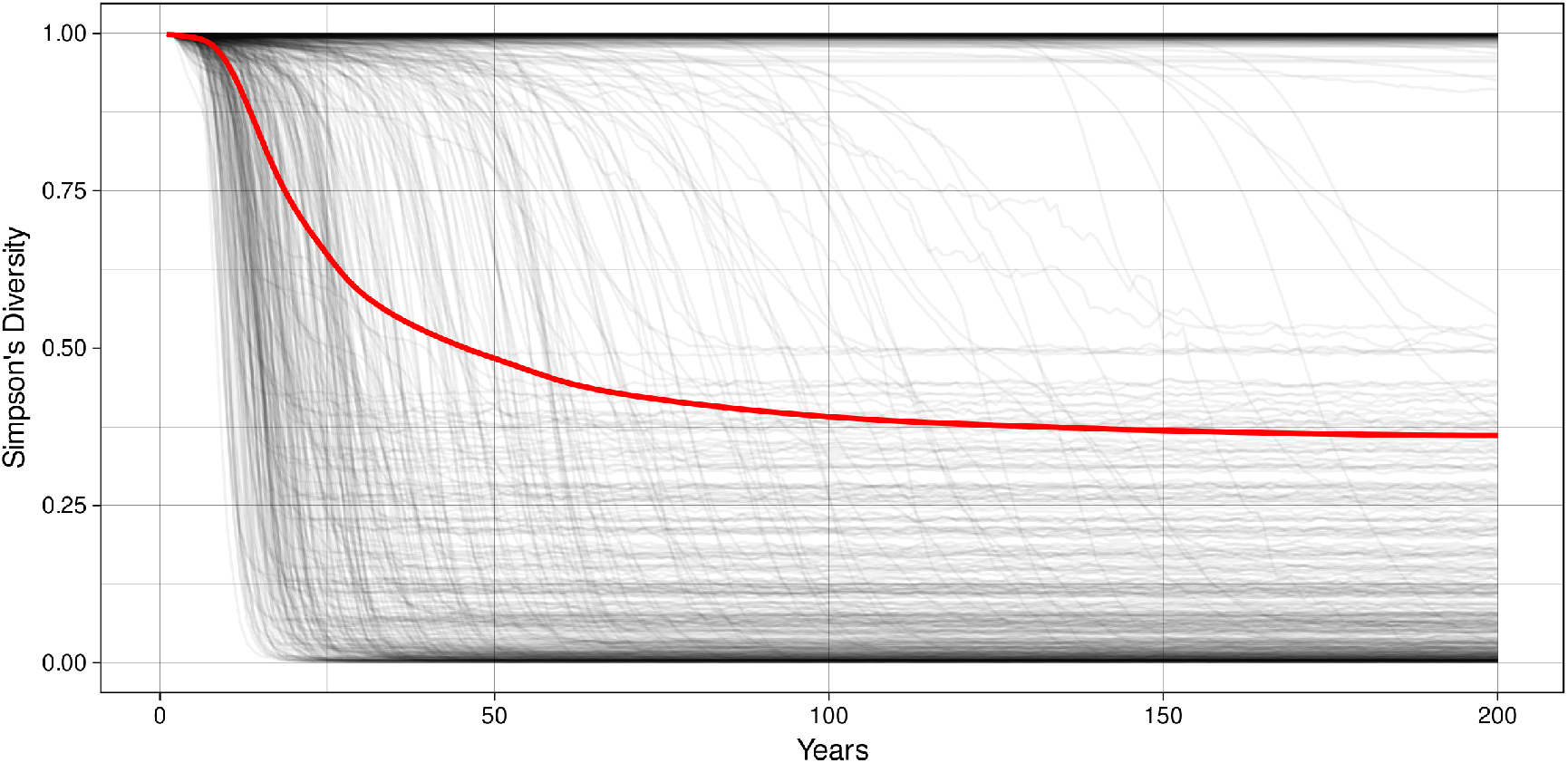
The Simpson’s diversity index for 500 burn-in iterations. The red line is the mean Simpson’s diversity index across all iterations in that year.

**Table S2:** The observed summary statistics from 1975, 2012, and 2019.

|  | *1975* | *2012* | *2019* |
| --- | --- | --- | --- |
| Proportion of syllables that appear once | 0.0094 | 0.0054 | 0.010 |
| Proportion of most common syllable type | 0.059 | 0.052 | 0.045 |
| Number of syllable types | 410 | 517 | 384 |
| Simpson's diversity index | 0.98 | 0.99 | 0.98 |
| Shannon's diversity index | 4.86 | 4.98 | 4.68 |
| Pielou's evenness index | 0.81 | 0.80 | 0.79 |
| Exponent of fitted power-law function | 9.17 | 108.81 | 1.96 |

**Figure S5:**
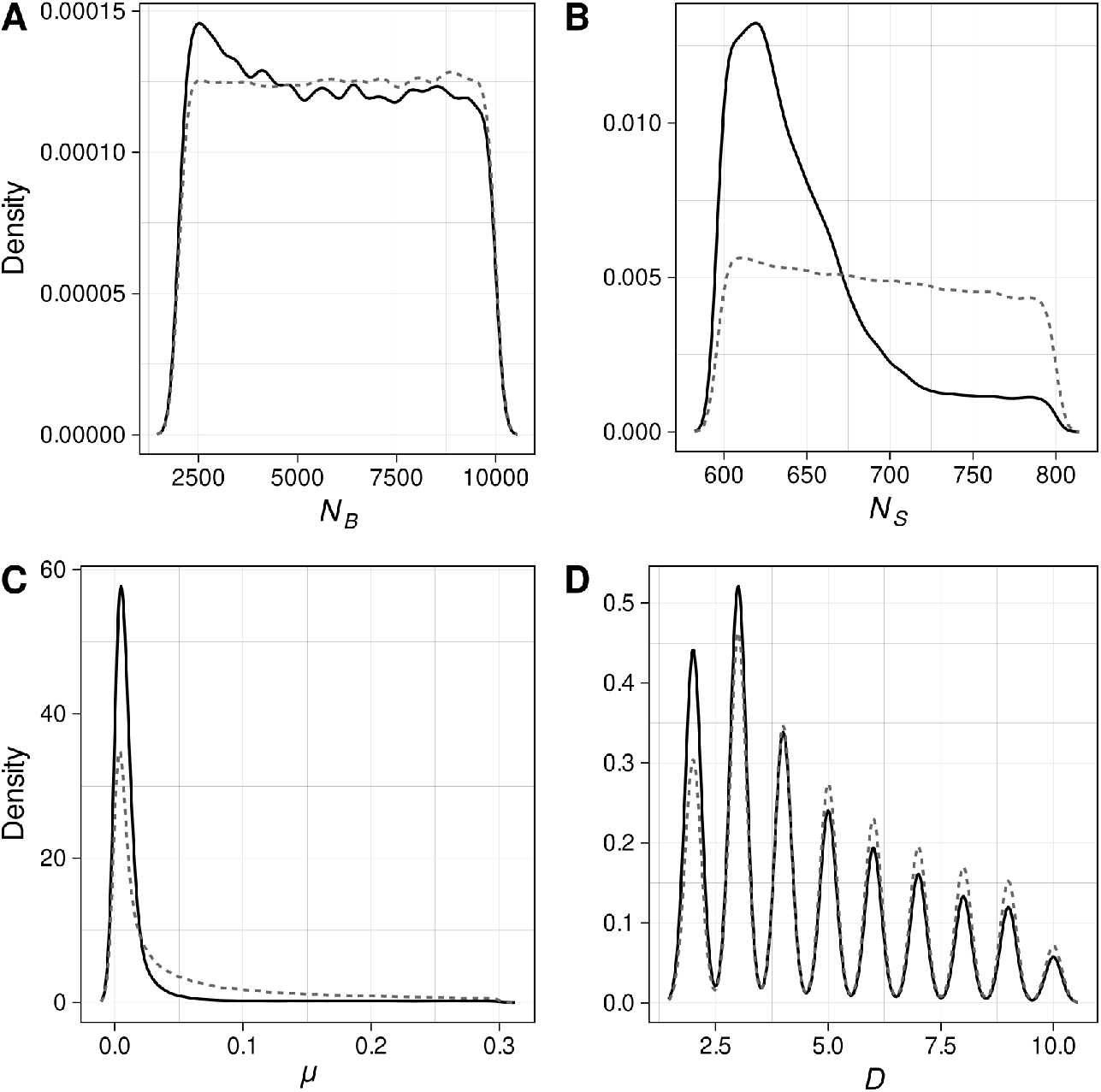
The prior (dotted lines) and posterior (solid lines) distributions for the four dynamic parameters in the agent-based model unrelated to transmission biases.

**Table S3:** The 10 most important summary statistics used by the random forests to estimate each parameter, identified using the Gini impurity method. The summary statistics are numbered as they appear in the methods: (1) the proportion of syllables that only appear once, (2) the proportion of the most common syllable type, (3) the number of syllable types, (4) Simpson’s diversity index, (5) Shannon’s diversity index, (6) Pielou’s evenness index, and (7) the exponent of the fitted power-law function to the progeny distribution.

|  | *1* | *2* | *3* | *4* | *5* | *6* | *7* | *8* | *9* | *10* |
| --- | --- | --- | --- | --- | --- | --- | --- | --- | --- | --- |
| $N_B$ | 2012 (6) | 2012 (2) | 2019 (6) | 1975 (6) | 2012 (4) | 1975 (4) | 1975 (5) | 2019 (2) | 2012 (1) | 2019 (4) |
| $N_S$ | 2012 (1) | 2012 (7) | 1975 (7) | 2012 (3) | 2019 (7) | 2019 (3) | 2019 (5) | 1975 (3) | 2012 (5) | 2012 (6) |
| $\mu$ | 2019 (1) | 1975 (1) | 2012 (3) | 2012 (1) | 2019 (3) | 1975 (3) | 2012 (5) | 2019 (5) | 2012 (6) | 1975 (6) |
| $D$ | 2019 (6) | 1975 (6) | 1975 (2) | 2019 (2) | 1975 (4) | 2012 (6) | 2012 (2) | 2012 (1) | 1975 (5) | 2019 (4) |
| $p_{att}$ | 1975 (5) | 2012 (5) | 1975 (4) | 2012 (4) | 2019 (2) | 2012 (2) | 2019 (5) | 1975 (3) | 2012 (1) | 2019 (4) |
| $\alpha$ | 1975 (2) | 2012 (2) | 1975 (4) | 2019 (2) | 2019 (4) | 2012 (4) | 2019 (5) | 1975 (5) | 2019 (6) | 2012 (5) |
| $v$ | 2019 (6) | 1975 (2) | 1975 (6) | 1975 (4) | 2019 (2) | 2012 (6) | 2012 (2) | 2012 (4) | 2012 (1) | 1975 (5) |

## References

1. Freeberg, T. M. Culture and courtship in vertebrates: a review of social learning and transmission of courtship systems and mating patterns. Behavioural Processes 51, 177–192 (2000).

2. Lipkind, D. & Tchernichovski, O. Quantification of developmental birdsong learning from the subsyllabic scale to cultural evolution. Proceedings of the National Academy of Sciences 108, 15572– 15579 (2011).

3. Mundinger, P. & Lahti, D. Quantitative integration of genetic factors in the learning and production of canary song. Proceedings of the Royal Society B 281, 20132631 (2014).

4. Aplin, L. M. Culture and cultural evolution in birds: a review of the evidence. Animal Behaviour 147, 179–187 (2019).

5. Slater, P. Bird song learning: causes and consequences. Ethology Ecology & Evolution 1, 19–46 (1989).

6. Parker, K. A., Anderson, M. J., Jenkins, P. F. & Brunton, D. H. The effects of translocation-induced isolation and fragmentation on the cultural evolution of bird song. Ecology Letters 15, 778– 785 (2012).

7. Cavalli-Sforza, L. L. & Feldman, M. W. Cultural Transmission and Evolution: A Quantitative Approach (Princeton University Press, Princeton, NJ, 1981).

8. Leonard, R. D. & Jones, G. T. Elements of an inclusive evolutionary model for archaeology. Journal of Anthropological Archaeology 6, 199–219 (1987).

9. Durham, W. H. Advances in Evolutionary Culture Theory. Annual Review of Anthropology 19, 187–210 (1990).

10. Boyd, R. & Richerson, P. J. in History and Evolution (eds Nitecki, M. H. & Nitecki, D. V.) 179–209 (State University of New York Press, Albany, NY, 1992).

11. Ormrod, R. K. Adaptation and Cultural Diffusion. Journal of Geography 91, 258–262 (1992).

12. Aplin, L. M. et al.. Experimentally induced innovations lead to persistent culture via conformity in wild birds. Nature 518, 538– 541 (2014).

13. Lachlan, R. F., Ratmann, O. & Nowicki, S. Cultural conformity generates extremely stable traditions in bird song. Nature Communications 9 (2018).

14. Neiman, F. D. Stylistic Variation in Evolutionary Perspective: Inferences from Decorative Diversity and Interassemblage Distance in Illinois Woodland Ceramic Assemblages. American Antiquity 60, 7–36 (1995).

15. Bentley, R. A. & Shennan, S. J. Cultural Transmission and Stochastic Network Growth. American Antiquity 68, 459–485 (2003).

16. Bentley, R. A., Lipo, C. P., Herzog, H. A. & Hahn, M. W. Regular rates of popular culture change reflect random copying. Evolution and Human Behavior 28, 151–158 (2007).

17. Youngblood, M. Conformity bias in the cultural transmission of music sampling traditions. Royal Society Open Science 6, 191149 (2019).

18. Byers, B. E., Belinsky, K. L. & Bentley, R. A. Independent cultural evolution of two song traditions in the chestnut-sided warbler. The American Naturalist 176, 476–489 (2010).

19. Kandler, A. & Powell, A. Generative inference for cultural evolution. Philosophical Transactions of the Royal Society B: Biological Sciences 373 (2018).

20. Crema, E. R., Edinborough, K., Kerig, T. & Shennan, S. J. An Approximate Bayesian Computation approach for inferring patterns of cultural evolutionary change. Journal of Archaeological Science 50, 160–170 (2014).

21. Kandler, A. & Crema, E. R. in Handbook of Evolutionary Research in Archaeology (ed Prentiss, A. M.) 83–108 (Springer International Publishing, Cham, 2019).

22. Crema, E. R., Kandler, A. & Shennan, S. Revealing patterns of cultural transmission from frequency data: Equilibrium and non-equilibrium assumptions. Scientific Reports 6, 1–10 (2016).

23. Kandler, A. & Shennan, S. A generative inference framework for analysing patterns of cultural change in sparse population data with evidence for fashion trends in LBK culture. Journal of the Royal Society Interface 12 (2015).

24. Raynal, L. et al.. ABC random forests for Bayesian parameter inference. Bioinformatics 35, 1720–1728 (2019).

25. Thompson, W. L. Agonistic behavior in the House Finch. Part I: Annual cycle and display patterns. The Condor 62, 245–271 (1960).

26. Mann, D. C., Lahti, D. C., Waddick, L. & Mundinger, P. C. House finches learn canary trills. Bioacoustics, 1–17 (2020).

27. Mundinger, P. C. Animal Cultures and a General Theory of Cultural Evolution. Ethology and Sociobiology 1, 183–223 (1980).

28. Nolan, P. M. & Hill, G. E. Female choice for song characteristics in the house finch. Animal Behaviour 67, 403–410 (2004).

29. Elliott, J. J. & Arbib, R. S. J. Origin and status of the House Finch in the eastern United States. Auk 70, 31–37 (1953).

30. Aldrich, J. W. & Weske, J. S. Origin and Evolution of the Eastern House Finch Population. The Auk 95, 528–536 (1978).

31. Belthoff, J. R. & Gauthreaux, S. A. J. Aggression and dominance in house finches. The Condor 93, 1010–1013 (1991).

32. Mundinger, P. C. Song dialects and colonization in the house finch, Carpodacus mexicanus, on the east coast. The Condor 77, 407– 422 (1975).

33. Ju, C., Geller, F. C., Mundinger, P. C. & Lahti, D. C. Four decades of cultural evolution in House Finch songs. The Auk: Ornithological Advances 136, 1–18 (2019).

34. Hawley, D. M., Hanley, D., Dhondt, A. A. & Lovette, I. J. Molecular evidence for a founder effect in invasive house finch (Carpodacus mexicanus) populations experiencing an emergent disease epidemic. Molecular Ecology 15, 263–275 (2006).

35. Boyd, R. & Richerson, P. J. Culture and the Evolutionary Process (The University of Chicago Press, Chicago, 1985).

36. Rendell, L. et al.. Cognitive culture: Theoretical and empirical insights into social learning strategies. Trends in Cognitive Sciences 15, 68–76 (2011).

37. Nelson, D. A., Hallberg, K. I. & Soha, J. A. Cultural evolution of puget sound white-crowned sparrow song dialects. Ethology 110, 879–908 (2004).

38. Soha, J. A. & Marler, P. A species-specific acoustic cue for selective song learning in the white-crowned sparrow. Animal Behaviour 60, 297–306 (2000).

39. Mennill, D. J., Badyaev, A. V., Jonart, L. M. & Hill, G. E. Male house finches with elaborate songs have higher reproductive performance. Ethology 112, 174–180 (2006).

40. Hernandez, A. M., Phillmore, L. S. & MacDougall-Shackleton, S. A. Effects of learning on song preferences and Zenk expression in female songbirds. Behavioural Processes 77, 278–284 (2008).

41. Jenkins, P. F. Cultural transmission of song patterns and dialect development in a free-living bird population. Animal Behaviour 25, 50–78 (1977).

42. Slater, P. J. B. & Ince, S. A. Cultural evolution in chaffinch song. Behaviour 71, 146–166 (1979).

43. Nelson, D. A. & Poesel, A. Does learning produce song conformity or novelty in white-crowned sparrows, Zonotrichia leucophrys? Animal Behaviour 78, 433–440 (2009).

44. Henrich, J. & Boyd, R. The Evolution of Conformist Transmission and the Emergence of Between-Group Differences. Evolution and Human Behavior 19, 215–241 (1998).

45. O’Loghlen, A. L. & Rothstein, S. I. Culturally correct song dialects are correlated with male age and female song preferences in wild populations of brown-headed cowbirds. Behavioral Ecology and Sociobiology 36, 251–259 (1995).

46. Searcy, W. A., Nowicki, S. & Hughes, M. The Response of Male and Female Song Sparrows to Geographic Variation in Song. The Condor 99, 651–657 (1997).

47. Eriksson, K., Enquist, M. & Ghirlanda, S. Critical points in current theory of conformist social learning. Journal of Evolutionary Psychology 5, 67–87 (2008).

48. Kandler, A. & Laland, K. N. An investigation of the relationship between innovation and cultural diversity. Theoretical Population Biology 76, 59–67 (2009).

49. Gibbs, H. L. Cultural evolution of male song types in Darwin’s medium ground finches, Geospiza fortis. Animal Behaviour 39, 253–263 (1990).

50. Garland, E. C. & McGregor, P. K. Cultural Transmission, Evolution, and Revolution in Vocal Displays: Insights From Bird and Whale Song. Frontiers in Psychology 11 (2020).

51. Otter, K. A., Mckenna, A., LaZerte, S. E. & Ramsay, S. M. Continent-wide Shifts in Song Dialects of White-Throated Sparrows. Current Biology 30, 3231–3235.e3 (2020).

52. Hernandez, A. M. & MacDougall-Shackleton, S. A. Effects of early song experience on song preferences and song control and auditory brain regions in female house finches (Carpodacus mexicanus). Journal of Neurobiology 59, 247–258 (2004).

53. Payne, R. B. Song learning and social interaction in indigo buntings. Animal Behaviour 29, 688–697 (1981).

54. Clayton, N. S. Song tutor choice in zebra finches. Animal Behaviour 35, 714–721 (1987).

55. Ciaburri, I. & Williams, H. Context-dependent variation of house finch song syntax. Animal Behaviour 147, 33–42 (2019).

56. Hoppitt, W. & Laland, K. N. in Advances in the Study of Behavior 105–160 (2008).

57. Barrett, B. J. Equifinality in empirical studies of cultural transmission. Behavioural Processes 161, 129–138 (2018).

58. Acerbi, A., Van Leeuwen, E. J., Haun, D. B. & Tennie, C. Conformity cannot be identified based on population-level signatures. Scientific Reports 6, 1–9 (2016).

59. Sardá-Espinosa, A. Time-Series Clustering in R Using the dtwclust Package. The R Journal (2019).

60. Ratanamahatana, C. A. & Keogh, E. Three myths about dynamic time warping data mining in Proceedings of the 2005 SIAM International Conference on Data Mining (2005), 506–510.

61. Roginek, E. W. Spatial variation of house finch (Haemorhous mexicanus) song along the American Southwest coast PhD thesis (Queens College, 2018), 1–51.

62. Müllner, D. fastcluster: Fast Hierarchical, Agglomerative Clustering Routines for R and Python. Journal of Statistical Software 53, 1–18 (2013).

63. Langfelder, P., Zhang, B. & Horvath, S. dynamicTreeCut: Methods for Detection of Clusters in Hierarchical Clustering Dendrograms R package version 1.63-1 (2016). https://CRAN.R-project.org/package=dynamicTreeCut.

64. Bitterbaum, E. & Baptista, L. F. Geographical variation in Songs of California House Finches (Carpodacus mexicanus). The Auk 96, 462–474 (1979).

65. Youngblood, M. A Raspberry Pi-based, RFID-equipped birdfeeder for the remote monitoring of wild bird populations. Ringing & Migration (2020).

66. Caldwell, C. A., Cornish, H. & Kandler, A. Identifying innovation in laboratory studies of cultural evolution: rates of retention and measures of adaptation. Philosophical Transactions of the Royal Society B 371 (2016).

67. Burnell, K. Cultural variation in savannah sparrow, Passerculus sandwichensis, songs: an analysis using the meme concept. Animal Behaviour 56, 995–1003 (1998).

68. Badyaev, A. V. & Martin, T. E. Sexual dimorphism in relation to current selection in the house finch. Evolution 54, 987–997 (2000).

69. Badyaev, A. V., Hill, G. E., Stoehr, A. M., Nolan, P. M. & McGraw, K. J. The evolution of sexual size dimorphism in the House Finch. II. Population divergence in relation to local selection. Evolution 54, 2134–2144 (2000).

70. Badyaev, A. V., Young, R. L., Oh, K. P. & Addison, C. Evolution on a local scale: developmental, functional, and genetic bases of divergence in bill form and associated changes in song structure between adjacent habitats. Evolution 62, 1951–1964 (2008).

71. Cutler, A., Cutler, D. R. & Stevens, J. R. in Ensemble Machine Learning: Methods and Applications (eds Zhang, C. & Ma, Y.) 157–176 (Springer, New York, 2012).

72. Blum, M. G. B., Nunes, M. A., Prangle, D. & Sisson, S. A. A Comparative Review of Dimension Reduction Methods in Approximate Bayesian Computation. Statistical Science 28, 189–208 (2013).

73. Pudlo, P. et al. Reliable ABC model choice via random forests. Bioinformatics 32, 859–866 (2015).

74. Veit, R. R. & Lewis, M. A. Dispersal, Population Growth, and the Allee Effect: Dynamics of the House Finch Invasion of Eastern North America. The American Naturalist 148, 255–274 (1996).

75. Lewis, M. A. in Spatial Ecology: The Role of Space in Population Dynamics and Interspecific Interactions (eds Tilman, D. & Kareiva, P.) (Princeton University Press, 1997).

76. Goodrich, B., Gabry, J., Ali, I. & Brilleman, S. rstanarm: Bayesian applied regression modeling via Stan R package version 2.19.3. 2020. https://mc-stan.org/rstanarm.

77. Podos, J. et al. A fine-scale, broadly applicable index of vocal performance: frequency excursion. Animal Behaviour 116, 203–212 (2016).

78. Lahti, D. C., Moseley, D. L. & Podos, J. A Tradeoff Between Performance and Accuracy in Bird Song Learning. Ethology 117, 802– 811 (2011).

79. Giraudeau, M. et al. Song characteristics track bill morphology along a gradient of urbanization in house finches (Haemorhous mexicanus). Front Zool 11, 83 (2014).

80. Luther, D. & Baptista, L. Urban noise and the cultural evolution of bird songs. Proceedings of the Royal Society B 277, 469–473 (2010).

81. Fernández-Juricic, E. et al. Microhabitat Selection and Singing Behavior Patterns of Male House Finches (Carpodacus mexicanus) in Urban Parks in a Heavily Urbanized Landscape in the Western U.S. Urban Habitats 3, 49–69 (2005).

82. Bermúdez-Cuamatzin, E., Ríos-Chelén, A. A., Gil, D. & Garcia, C. M. Strategies of song adaptation to urban noise in the house finch: syllable pitch plasticity or differential syllable use? Behaviour 146, 1269–1286 (2009).

83. Bermúdez-Cuamatzin, E., Ríos-Chelén, A. A., Gil, D. & Garcia, C. M. Experimental evidence for real-time song frequency shift in response to urban noise in a passerine bird. Biol Lett 7, 36–38 (2011).

84. Brakes, P. et al. Animal cultures matter for conservation. Science 363, 1032–1034 (2019).

85. Caro, T. & Sherman, P. W. Vanishing behaviors. Conservation Letters 5, 159–166 (2012).

86. Keith, S. A. & Bull, J. W. Animal culture impacts species’ capacity to realise climate-driven range shifts. Ecography 40, 296–304 (2017).

87. Laiolo, P. & Jovani, R. The emergence of animal culture conservation. Trends in Ecology and Evolution 22, 5 (2007).

88. Whitehead, H. Conserving and managing animals that learn socially and share cultures. Learning and Behavior 38, 329–336 (2010).

89. United Nations Convention on Migratory Species. Decisions 12.75 to 12.77 - Conservation implications of animal culture and social complexity (2017).

90. Holland, J., McGregor, P. K. & Rowe, C. L. Changes in Microgeographic Song Variation of the Corn Bunting Miliaria calandra. Journal of Avian Biology 27, 47 (1996).

91. Briefer, E., Osiejuk, T. S., Rybak, F. & Aubin, T. Are bird song complexity and song sharing shaped by habitat structure? An information theory and statistical approach. Journal of Theoretical Biology 262, 151–164 (2010).

92. Hill, S. D. & Pawley, M. D. Reduced song complexity in founder populations of a widely distributed songbird. Ibis 161, 435–440 (2019).

93. Laiolo, P. & Tella, J. L. Habitat fragmentation affects culture transmission: patterns of song matching in Dupont’s lark. Journal of Applied Ecology 42, 1183–1193 (2005).

94. Laiolo, P. & Tella, J. L. Erosion of animal cultures in fragmented landscapes. Frontiers in Ecology and the Environment 5, 68–72 (2007).

95. Ortega, Y. K., Benson, A. & Greene, E. Invasive plant erodes local song diversity in a migratory passerine. Ecology 95, 458–465 (2014).

96. Pang-Ching, J. M., Paxton, K. L., Paxton, E. H., Pack, A. A. & Hart, P. J. The effect of isolation, fragmentation, and population bottlenecks on song structure of a Hawaiian honeycreeper. Ecology and Evolution 8, 2076–2087 (2018).

97. Paxton, K. L. et al. Loss of cultural song diversity and the convergence of songs in a declining Hawaiian forest bird community. Royal Society Open Science 6, 190719 (2019).

98. Petrusková, T., Osiejuk, T. S. & Petrusek, A. Geographic variation in songs of the tree pipit (Anthus trivialis) at two spatial scales. The Auk 127, 274–282 (2010).

99. Sebastián-González, E. & Hart, P. J. Birdsong meme diversity in a habitat landscape depends on landscape and species characteristics. Oikos 126, 1511–1521 (2017).

100. Laiolo, P., Vögeli, M., Serrano, D. & Tella, J. L. Song Diversity Predicts the Viability of Fragmented Bird Populations. PLOS ONE 3, e1822 (2008).

101. Laiolo, P. Characterizing the spatial structure of songbird cultures. Ecological Applications 18, 1774–1780 (2008).

102. Pérez-Granados, C., Osiejuk, T. & López-Iborra, G. M. Habitat fragmentation effects and variations in repertoire size and degree of song sharing among close Dupont’s Lark Chersophilus duponti populations. Journal of Ornithology 157, 471–482 (2016).

103. Garland, E. C. et al. Population structure of humpback whales in the western and central South Pacific Ocean as determined by vocal exchange among populations. Conservation Biology 29, 1198–1207 (2015).

104. Rivera-Gutierrez, H. F., Matthysen, E., Adriaensen, F. & Slabbekoorn, H. Repertoire sharing and song similarity between great tit males decline with distance between forest fragments. Ethology 116, 951–960 (2010).

105. Greggor, A. L., Thornton, A. & Clayton, N. S. Harnessing learning biases is essential for applying social learning in conservation. Behavioral Ecology and Sociobiology 71 (2017).

106. Barrett, B., Zepeda, E., Pollack, L., Munson, A. & Sih, A. Counterculture: Does social learning help or hinder adaptive response to human-induced rapid environmental change? Frontiers in Ecology and Evolution 7, 1–18 (2019).

## References

1. Brock, G., Pihur, V., Datta, S. & Datta, S. clValid: An R Package for Cluster Validation. Journal of Statistical Software 25 (2008).

2. Arbelaitz, O., Gurrutxaga, I., Muguerza, J., Pérez, J. M. & Perona, I. An extensive comparative study of cluster validity indices. Pattern Recognition 46, 243–256 (2013).

